# Extracting the phylogenetic dimension of coevolution reveals hidden functional signal

**DOI:** 10.1101/2020.09.23.310300

**Authors:** Alexandre Colavin, Esha Atolia, Anne-Florence Bitbol, Kerwyn Casey Huang

**Author notes:** Co-first authors (Alexandre Colavin, Esha Atolia).

## Abstract

Despite the structural and functional information contained in the statistical coupling between pairs of residues in a protein, coevolution associated with function is often obscured by artifactual signals such as genetic drift, which shapes a protein’s phylogenetic history and gives rise to concurrent variation between protein sequences that is not driven by selection for function. Here, we introduce a method for explicitly defining a phylogenetic dimension of coevolution signal, and demonstrate that coevolution can occur on multiple phylogenetic timescales within a single protein. Our method, Nested Coevolution (NC), can be applied as an extension to any coevolution metric. We use NC to demonstrate that poorly conserved residues can nonetheless have important roles in protein function. Moreover, NC improved structural-contact prediction over gold-standard coevolution-based methods, particularly in subsampled alignments with fewer sequences. NC also lowered the noise in detecting functional sectors of collectively coevolving residues. Sectors of coevolving residues identified after NC correction were more spatially compact and phylogenetically distinct from the rest of the protein, and strongly enriched for mutations that disrupt protein activity. Our conceptualization of the phylogenetic separation of coevolution represents an advance from previous pragmatic attempts to reduce phylogenetic artifacts in measurements of coevolution. Application of NC broadens the application of protein coevolution measurements, particularly to eukaryotic proteins with fewer naturally available sequences, and further elucidates relationships among protein evolution and genetic diseases.

## Introduction

It has long been appreciated that comparisons among homologous sequences of a protein of interest can provide key information about its function and structure. Just as evolutionarily conserved individual residues are generally crucial to a protein’s proper function, the statistical covariation (arising from correlated evolution, i.e. coevolution) between pairs of residues (1, 2) carries information that is useful for predicting structural contacts (3–7) and protein-protein interactions (8–11) and their interfaces (12), intuiting novel protein conformations (5), understanding protein allostery (13), interpreting variants (14), identifying functional domains (15–18), and reprograming protein specificity (19). However, despite the increasing prevalence of sequencing data, sampling of the phylogenetic tree is necessarily limited and biased. Evolutionary events such as speciation can drive simultaneous changes that are statistically linked but may not reflect relevant functional coupling, for example when they arise from genetic drift. Hence, spurious covariation is more likely to arise in comparisons between distantly related sequences, hindering the ability of such studies to deliver functional insights.

Of the numerous existing methods for measuring protein coevolution, many implement methods for reducing the effects of phylogenetic noise. Although mutual information is extremely sensitive to the phylogenetic distribution of sequences and the conservation (measured via entropy) of individual positions, normalization by the joint entropy reduces the influence of phylogeny and entropy and improves structural-contact prediction (20). Statistical coupling analysis, which normalizes the covariance matrix by a function of the entropy, provides sufficient information to specify a protein fold (21) and to detect functional domains (6, 18). Direct coupling analysis (DCA) usually involves down-weighting the coevolutionary signal contributions from over-represented sequences, and attempts to deconvolve higher-order correlations to identify directly interacting residue pairs (4, 22). Motivated by the observed strong relationship between a position’s average mutual information and the mutual information it exhibits with specific positions, modifications such as the average product correction (APC) subtract this average signal; this correction can be applied to any existing coevolution metric other than mutual information. However, none of these strategies attempt to resolve the evolutionary timescale of coevolution.

Even with affordable sequencing and widespread environmental sampling, coevolution methods are often limited by the number of naturally occurring protein sequences available. Successful predictions of structural contacts often require several thousand sequences to align (3, 23), which is generally prohibitive for many mammalian proteins. For other proteins, the phylogenetic distribution of available sequences is skewed by sampling and is well recognized as a source of spurious signal in coevolution (20, 24). Thus, methods that enable the separation of functional coupling from phylogenetic and sampling noise would greatly expand the utility of coevolution, particularly for applications to diseases involving human proteins with limited numbers of available sequences.

Here, we introduce the concept of Nested Coevolution (NC), a correction that leverages a well-defined null hypothesis to accurately measure the coevolutionary signal above what is expected from phylogenetic distribution alone. We determined that NC results in higher fidelity of the coevolutionary signal across gold-standard coevolution-based metrics for structural prediction for many proteins, especially with fewer sequences. In addition, we found that NC improves the detection of spatially contiguous groups of collectively coevolving residues (“sectors”) that are phylogenetically distinct from each other and the protein itself, beyond differences in entropy alone. Finally, sectors identified using NC were enriched for positions at which mutations are maximally deleterious, highlighting the functional significance of signal from our method. Since our method is agnostic to the underlying method of measuring coevolution, we anticipate wide utility for the ability to resolve the temporal dimension of protein coevolution.

## Results

### Background model of coevolution reveals temporal dimension of coevolution

To interrogate the contribution of phylogenetic sampling to protein coevolution measurements, we sought to separate the coevolution signal due to inter-clade and intra-clade sequence comparisons (Fig. 1A,B). Given a multiple sequence alignment (MSA) for a protein of interest (Fig. 1Ai), we first measure the total covariation (*C*_*T*_) between every pair of positions (Fig. 1Aii) using an established metric of residue-residue coupling such as the normalized mutual information (NMI; Fig. 1A) (20):

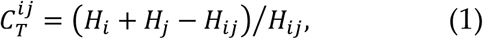

where *H*_*i*_ is the Shannon entropy (a measure of conservation) of position *i*, and *H*_*ij*_ is the joint Shannon entropy of positions *i* and *j*. The quantity *H*_*i*_ + *H*_*j*_ − *H*_*ij*_ is the mutual information between positions *i* and *j*, which measures the coupling between residues (Fig. S1). The NMI residue pair covariation in Eq. 1 is a natural metric choice because normalizing by *H*_*ij*_ makes the mutual information independent of conservation (20). Note that our algorithm can be applied to any covariation metric, and as we will show, our main results are robust to metric choice.

**Figure 1:**
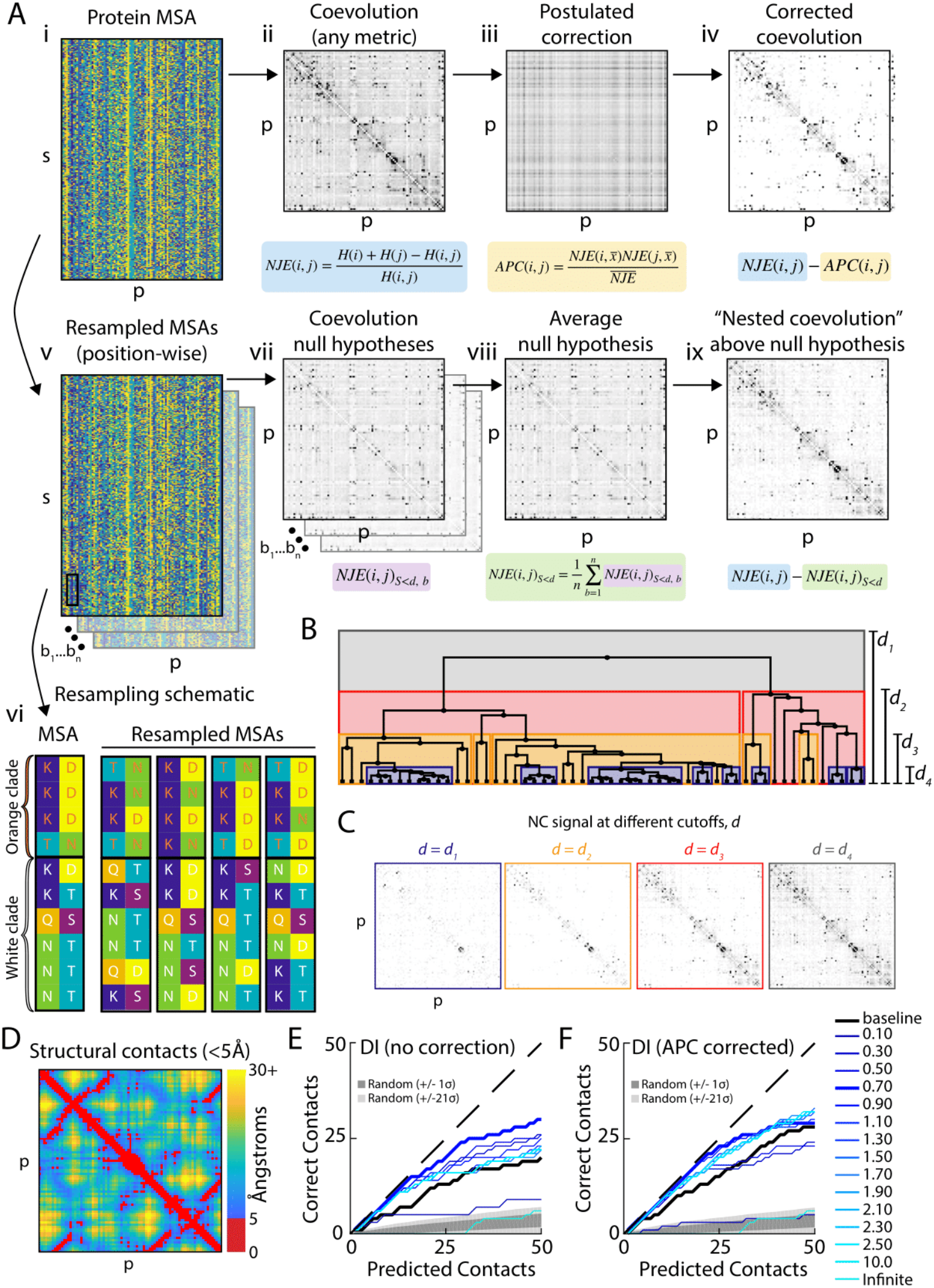
NC introduces a phylogenetic dimension to traditional coevolution metrics that removes noise and improves structural prediction. A) Schematic illustrating the NC correction to traditional coevolution algorithms. The MSA (i) is used to generate a covariation matrix (ii) with a particular metric such as normalized joint entropy or direct information. Previous studies have attempted to remove phylogenetic noise using the APC (iii), which results in a corrected coevolution matrix (iv) that has lower levels of off-diagonal signal. For the NC correction, the MSA is resampled multiple times (v) within clades defined by a phylogenetic cutoff *d* (vi), providing null hypotheses (vii) that are averaged (viii) to correct the covariation matrix (i). The resulting difference (ix) is the NC matrix for a particular cutoff *d*. B) The Jukes-Cantor phylogenetic distance between homologs defines clades (visualized as a tree) within the NC cutoff *d*. C) NC signal at different cutoffs *d* as illustrated in (B) for the MSA of the KH domain from (B). For small values of *d*, the NC matrix exhibits very little off-diagonal signal, signifying a reduction in noise. D) The structural contact map for KH, highlighting contacts that are in close 3D proximity (<5 Å, red), respectively. E,F) NC with particular cutoffs *d* improves the prediction of structural contacts relative to DCA, applied to DI without (E) or after correction with APC (F) (black lines). All residues within five positions on the polypeptide sequence were excluded from the analysis. Gray represents the predictions of the baseline NMI metric.

The most straightforward null hypothesis for protein coevolution is that coevolutionary coupling between pairs of proteins is completely absent—that is, that the probability of a position having any particular amino acid identity is independent of any other position’s identity. Although this null hypothesis can be evaluated analytically for some methods (SI), other methods have no known closed-form solution for the expected value of the coevolution matrix under these conditions. Hence, we computationally compute the average coevolution signal from many globally resampled MSAs in which each position in each protein in the original MSA is replaced by the equivalent position from another randomly chosen protein (resampled with replacement; Fig. 1Av,vi). We expect any measured coevolution from these resampled matrices to represent signal due simply to the distribution of amino acid identities at each position; any significant difference between the coevolution signal measured in the original MSA and this null hypothesis can potentially be attributed to coevolution.

However, this initial null hypothesis does not test for the phylogenetic structure of sequences; in the globally resampled MSAs, every sequence is effectively evolutionarily equidistant from one another. Previous attempts to remove the influence of phylogeny such as APC (Fig. 1Aiii), which corrects the covariation matrix by subtracting the product of its mean value across columns and rows for each pair of positions (Fig. 1Aiv), have substantially improved contact prediction (20). However, the APC is a postulated correction that does not directly take into account the phylogenetic structure of an MSA. We sought to construct a null hypothesis-driven background model of the expected coevolution in an MSA in which intra-clade coevolution is explicitly removed. We achieve this goal by generating MSAs by resampling each position from sequences that are closely related (Fig. 1Av,vi), thus removing correlations arising from recent evolutionary history within each clade. We define a clade as the subset of sequences *S* with a Jukes-Cantor distance below *d*, which we refer to as the phylogenetic cutoff. For each value of *d*, we calculate the inter-clade covariation 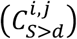 from a resampled MSA either analytically or via bootstrapping (Fig. 1Avii, S2A, Methods), where *C* denotes the chosen covariation measure (e.g. NMI). This inter-clade covariation thus measures the expected value of covariation due solely to the comparison of sequences between clades (Fig. 1B). We then average over many such null hypotheses (over many within-clade resampled MSAs at fixed *d*), yielding the mean inter-clade covariation matrix (*C*_*S*>*d*_) (Fig. 1Aviii), which represents the expected coevolution due to both the distribution of amino acid identities at each position and the phylogenetic structure of the protein MSA (Fig. 1B). Significant differences between this background model and the baseline signal measured from the original MSA represent signal that was contained in the intra-clade comparison of closely related sequences. Since the difference between the background model and the baseline signal qualitatively captures the significance of the baseline measurement (Fig S2B, Methods), we subtract *C*_*S*>*d*_ from the total covariation *C*_*T*_ to obtain the phylogenetic cutoff-dependent covariation signal *C*_*S*≤*d*_ (Fig. 1Aix); positive values indicate that the total covariation is larger than expected by comparison of sequences between clades, thus revealing covariation arising from recent evolutionary history in all clades:

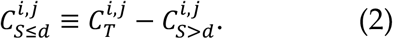

We refer to the signal 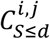 above the null hypothesis 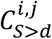 in Eq. 2 as a protein’s “nested coevolution” (NC), in that it separates coevolution signal into signal attributed to comparison of sequences either within 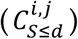 or between 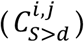 nested clades of a phylogenetic tree. The only free parameter in the NC is the phylogenetic cutoff (*d*). As we vary the cutoff value, many patterns of NC typically emerge, revealing distinct windows of coevolution for a single protein MSA (Fig. 1C). The changes in NC observed between two cutoffs represent the signal due to pairs of sequences whose distance is between the cutoffs used to calculate each window. Hence, distinct evolutionary timescales of protein coevolution are revealed as the phylogenetic cutoff is varied.

To test the relevance of NC windows to protein structure prediction, we measured the enrichment of structural contacts from the pairs of residues with the highest 50 values in the NC matrix 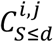 for each value of *d*. Here, we applied NC as a correction to DCA, the current gold standard for coevolution-based prediction of structural contacts (4, 22). We employed the direct information (DI) metric to quantify coevolution (4, 22). In this and subsequent analyses, we considered structural contacts to be within 5 Å at closest approach, excluding pairs of residues within 5 amino acids on the sequence (Methods).

The NC phylogenetic cutoffs revealed a variety of improvements for the KH domain (Fig. 1D), which is present in a wide variety of nucleic acid-binding proteins (25). Some windows generally outperformed DCA, without (Fig. 1E) or with (Fig. 1F) the APC.

To determine the added value of NC for other proteins and for another frequently utilized coevolution metric, the Frobenius norm (26), which is frequently utilized in DCA as an alternative to DI (27, 28), we carried out a DCA structural-contact analysis for 10 protein family domains with DI or Frobenius norm (Methods). Across both metrics and all proteins, NC improved the predictions of structural contacts (Fig. 2A), even relative to the inclusion of APC (20). Hence, NC is a correction that enhances the predictive power of state-of-the-art coevolution measurements.

**Figure 2:**
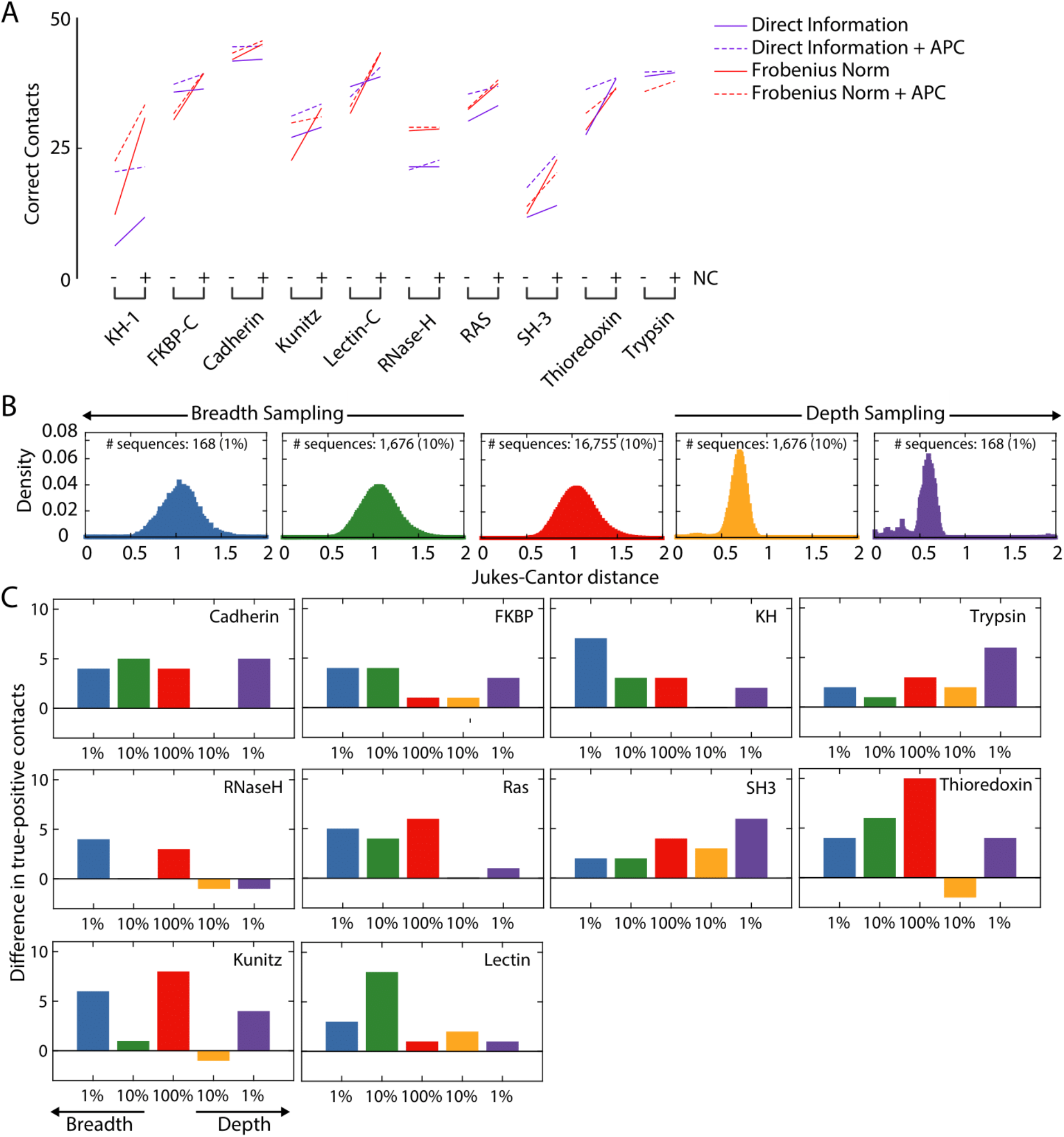
NC improves predictions of structural contacts across proteins and coevolution methods, and resolves information loss due to subsampling of the set of sequences. A) NC increased the number of true-positive structural contacts among the first 50 predictions for 10 highly conserved proteins predicted by DCA using DI or Frobenius norm, without or with APC. B) MSAs were subsampled across breadth (random sampling) and depth (sorted sampling) of the MSA. Typically, the distribution of Jukes-Cantor distances in the MSA (red) remained essentially unchanged for breadth sampling (green and blue), while it shifted to lower values (as expected) for depth sampling (gold and purple); shown is the KH domain. C) NC generally increased the number of true-positive structural contacts among the first 50 predictions relative to DCA employing DI (without APC) across proteins and both breadth and depth sampling (for DI with APC, see Fig. S3). Small decreases occurred for depth sampling of RNase H, thioredoxin, and Kunitz.

### NC improves predictions of structural contacts using fewer sequences

One common limitation for computing coevolution is the number of homologous sequences available for constructing an MSA. To interrogate whether NC could still accurately predict structural contacts with fewer sequences, we subsampled the MSAs of 10 proteins with different breadth (randomly selecting 10% or 1% of the sequences) or depth (selecting the 10% or 1% of sequences most related to the protein used to construct the MSA, Table S1) (Fig. 2B). NC improved structural contact prediction for a majority of the subsampled MSAs when correcting DI without (Fig. 2C, S4A) or with (Fig. S3B) application of APC. For the KH domain, more than twice as many true positives were predicted after applying NC compared with DI+APC alone (Fig. S3A). Perhaps unsurprisingly, breadth sampling generally performed better than depth sampling (Fig. 2C), indicating that accurate prediction is reliant on the sequences being sufficiently distantly related. Nonetheless, for many proteins, the value of the NC correction was enhanced when the number of homologous sequences was low, both for depth and breadth samplings.

### NC generates eigenvectors with increased fidelity, improving detection of spatially contiguous sets of coevolving residues

Previous studies have utilized coevolution measurements to identify groups of residues within a protein that are spatially contiguous on the tertiary structure and thus are postulated to have a joint function (6, 18, 29–32). These “sectors” can by defined by a variety of methods, such as the extreme-value residues of the eigenvectors of the coevolution matrix with the largest eigenvalues (18), and have been proposed to reflect independent biological properties such as catalytic efficiency and thermal stability (18). Motivated by these successes, we sought to measure the effect of incorporating the phylogenetic dimension revealed by NC when defining sectors of residues. Specifically, we measured the NC- and APC-corrected coevolution using NMI across a range of phylogenetic cutoffs, concatenating the results and performing eigendecomposition to identify the most significant eigenvectors (Methods). The residues most strongly associated with the positive or negative components of each resulting eigenvector are considered a sector.

We first focused on MreB, an essential protein involved in cell-shape determination in many rod-shaped bacteria (33). MreB belongs to a protein family that includes ParM, FtsA, and MamK in bacteria, crenactin in archea, and actin in eukaryotes (34, 35). These proteins are structural homologs characterized by a four-subdomain fold around an ATP-binding pocket (35, 36), with very low sequence identity and disparate cellular functions. Thus, we anticipated that the set of MreB homologs would have sufficient diversity to support robust coevolution measurements, particularly functional sectors.

We compared NC-derived sectors with baseline sectors derived from eigenvectors of the baseline coevolution matrix for MreB homologs. We identified the most closely related baseline sectors for three NC eigenvectors with some of the highest eigenvalues, which we refer to as eigenvectors A, B, and C (Methods). Each pair of NC and baseline eigenvectors appeared similar, especially for the residues with the largest absolute coefficients (Fig. 3A-C). However, the baseline eigenvectors exhibited much higher variation of coefficients for residues across the protein (Fig. 3A-C). For eigenvectors A and B, the NC-derived eigenvectors exhibited 32.8-fold and 38.3-fold lower standard deviation (after removing the 50 highest and lowest coefficients) than the baseline-derived eigenvectors, respectively (Fig. 3A,B). For eigenvector C, the baseline eigenvector contained residues with both highly positive and highly negative coefficients, while the high-magnitude coefficients of the NC eigenvector were solely positive (Fig. 3C); the positive portion of the NC eigenvector again had substantially lower noise than the baseline eigenvector (2.1-fold lower standard deviation, Fig. 3C).

**Figure 3:**
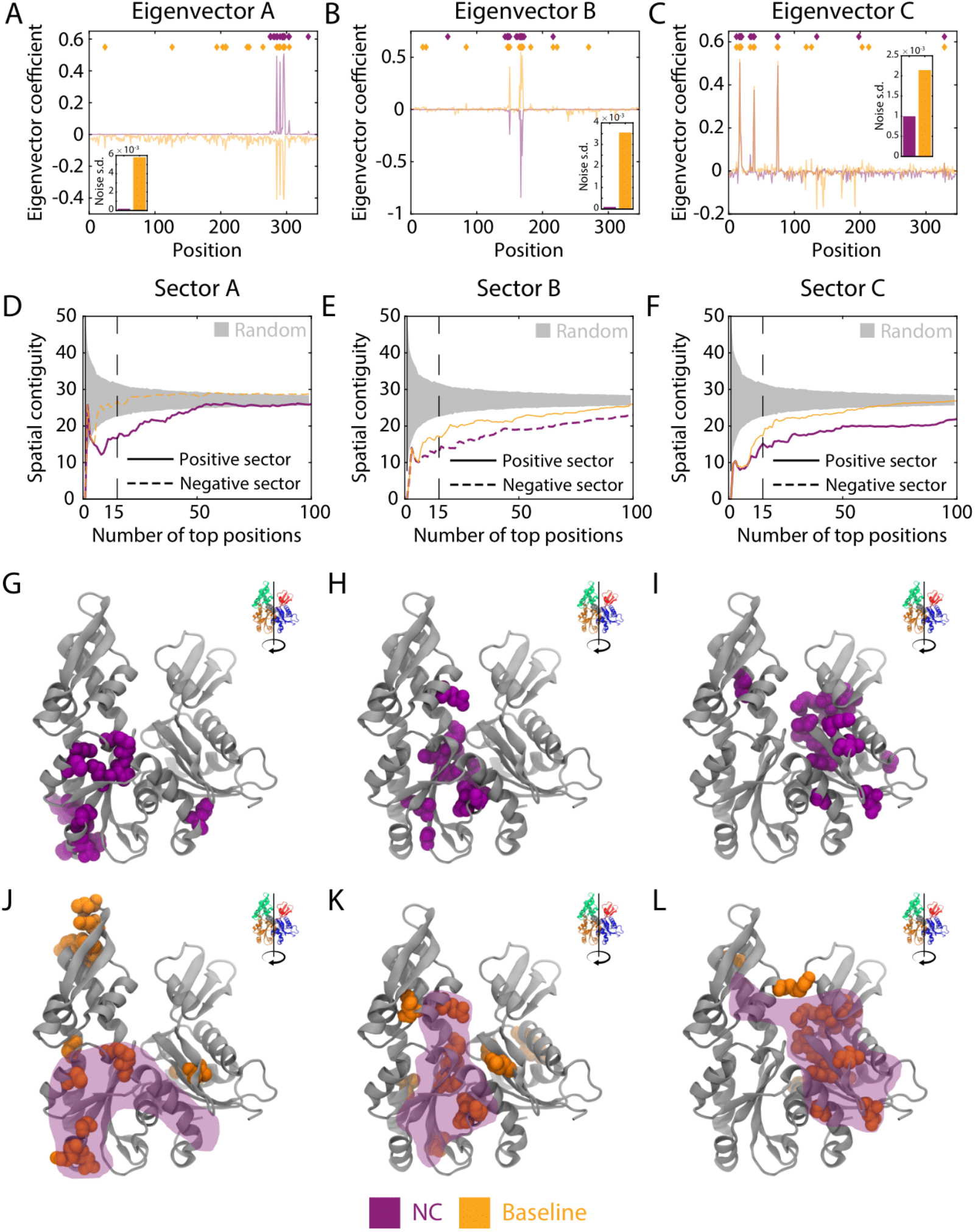
NC eigenvectors for the actin homolog MreB have lower noise and are more spatially contiguous than baseline eigenvectors. A-C) Three eigenvectors with large eigenvalues were identified and paired between baseline coevolution (NMI with APC) and the NC correction for an MSA containing 9,998 sequences of MreB. Aside from the residues with large coefficients, the NC eigenvectors exhibited lower background noise than the baseline eigenvectors. Insets: standard deviations of the eigenvector coefficients after excluding the highest and lowest 50 values. D-F) NC sectors are more spatially contiguous than the corresponding baseline sectors. Sectors were defined based on a sliding cutoff of the most positive or most negative coefficients of each eigenvector in (A-C). Spatial contiguity was defined as the mean pairwise distance between each residue within a sector. G-L) For the 15-residue versions of the NC and baseline sectors (vertical lines in (D-F)), the NC sectors (G-I) are more compact on the three-dimensional structure than the corresponding baseline sectors (J-L). The shaded purple regions in (J-L) represent the NC sector.

Motivated by the distinct behaviors of the positive and negative components of eigenvector C, we defined distinct positive and negative sectors (Methods) for each NC and baseline eigenvector using a variable cutoff on the site contributions to adjust sector size (as sectors are defined as the sets of amino acids with highest site contributions in a given eigenvector). For different sector sizes, we quantified the spatial contiguity as the mean pairwise distance between each residue within a sector. For sectors A-C (derived from eigenvectors A-C), the first 5-9 residues exhibited approximately the same spatial contiguity in the NC as in the baseline eigenvectors (Fig. 3D-F). However, as the cutoff was increased, the NC sector remained more spatially compact than the baseline sector (Fig. 3D-F). All three NC sectors were also more spatially contiguous than expected based on random sampling for cutoffs yielding at least 50 residues (Fig. 3D-F), while the baseline sector A was distributed across the protein structure (Fig. 3D,J). NC sectors A and C were largely situated in subdomains IIA (Fig. 3G) and IA (Fig. 3I), respectively, while sector B was localized to the ATP-binding pocket (Fig. 3H). Notably, sector C was spatially contiguous (Fig. 3I) despite being spread across the protein sequence (Fig. 3C). Baseline sectors B and C with 15 residues were qualitatively similar to the corresponding NC sectors (Fig. 3K,L); the large background fluctuations of the baseline eigenvector likely led to the inclusion of additional, erroneous residues into the sector prediction. Thus, the phylogenetic correction of NC improves the fidelity of sector detection as measured by the spatial contiguity of its constituent residues.

### Sectors display distinct phylogenetic signatures from the rest of the protein

Since sectors have been postulated to reflect distinct evolutionary histories driven by selection for particular biological functions (18), we sought to compare the phylogeny of the residues within a sector with other sectors and the rest of the protein. The MirrorTree algorithm (Methods) was originally developed to compare phylogenies of two proteins, motivated by the assumption that similar histories signifies a common function, e.g. through protein-protein interactions and/or acting in the same pathway (37, 38). After computing a pairwise distance matrix of all sequences within an MSA for each of the two proteins based on homologs in the same set of organisms, the MirrorTree score is defined as the Pearson correlation coefficient between the entries in the two pairwise distance matrices (37). We straightforwardly modified the MirrorTree method to compare the complete protein MSA to the MSA filtered to include only the residues within the sector of interest (Fig. 4A).

**Figure 4:**
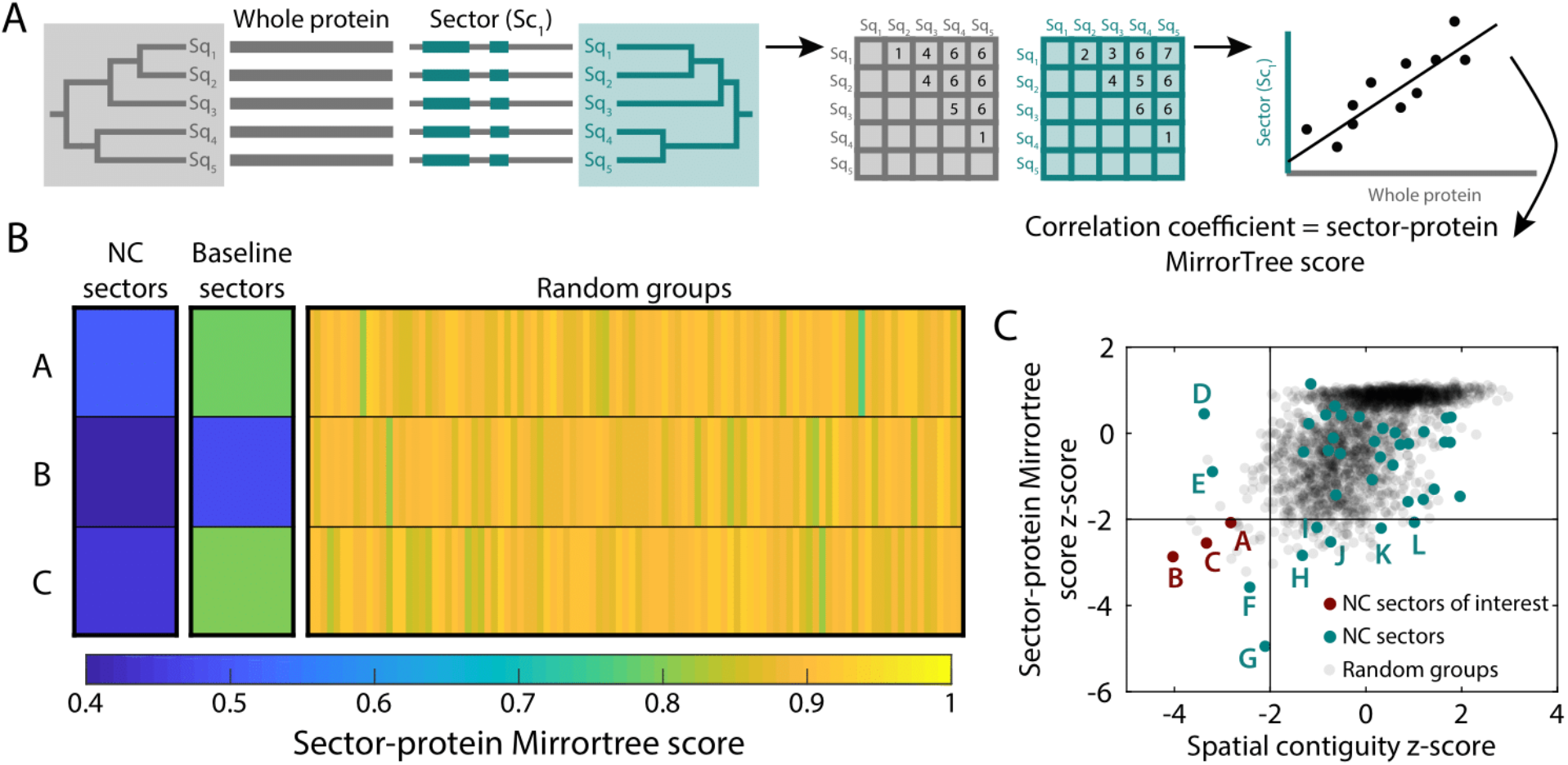
Sectors are phylogenetically distinct from the entire protein. A) Repurposing the MirrorTree algorithm (37) to measure the phylogenetic similarity between sectors and the entire protein. The MirrorTree score is defined as the Pearson correlation coefficient between the entries in the two pairwise distance matrices of all sequences within an MSA for the protein versus only the residues in the sector. B) MreB NC sectors A-C (Fig. 3) had lower sector-protein MirrorTree scores than the corresponding baseline sectors, while random groups of 15 residues had MirrorTree scores close to 1 (as expected). C) MreB NC sectors were computed from the 15 most positive or negative coefficients of the 20 eigenvectors with the highest eigenvalues. Among these 40 sectors, the *z*-scores of the MirrorTree score and the spatial contiguity were <-2 for sectors A-C. Sectors D-L substantially overlapped sectors A-C, and are considered in Fig. 6.

To broadly investigate sector identification, we identified 40 15-residue sectors for MreB based on the positive and negative coefficients of the 20 eigenvectors with the highest eigenvalues. As negative controls, we randomly sampled sets of residues of the same size as each sector from across the protein. Sector-protein MirrorTree scores for sectors A-C (Fig. 3) were substantially lower for sectors than for the random groups (Fig. 4B), which all had MirrorTree scores close to 1, as expected (Fig. 4B). Baseline sectors A-C had MirrorTree scores intermediate between those of the corresponding NC sector and random groups (Fig. 4B), likely reflecting the noisy selection from baseline eigenvectors of residues that functionally follow the phylogenetic history of the protein overall. To evaluate the significance of the MirrorTree score and of the spatial contiguity of each sector, we computed *z*-scores based on the mean and standard deviation of the two metrics applied to the random groups of the same size as each sector. Sectors A-C had MirrorTree scores <0.5 (Fig. 4B), indicating distinct phylogenetic histories from the protein, and MirrorTree and spatial contiguity *z*-scores<-2 (Fig. 4C). There were four other sectors (D-G) that had spatial contiguity *z*-scores<-2. These sectors largely overlapped with A-C; we will return to this overlap in a later section. All other sectors had spatial contiguity *z*-score>2, and all but five (H-L) had MirrorTree *z*-score>−2. Thus, MirrorTree reveals that certain NC sectors have distinct evolutionary trajectories from the protein itself, motivating us to focus on certain sectors (such as A, B, and C for MreB).

### Phylogenetic similarity and the role of entropy

Conservation itself is a major determinant of protein function (39–41), and spatially contiguous sets of residues can be identified solely on the basis of conservation (42). To account for variation in entropy across a protein, previous studies have excluded positions with high conservation (Shannon entropy<0.1) or composed of >25% gaps in the MSA (43). For MreB, NC sectors A-C had lower entropy than baseline sectors or random groups of the same size (Fig. 5A-C), albeit higher entropy than residues typically considered highly conserved (entropy<0.1).

**Figure 5:**
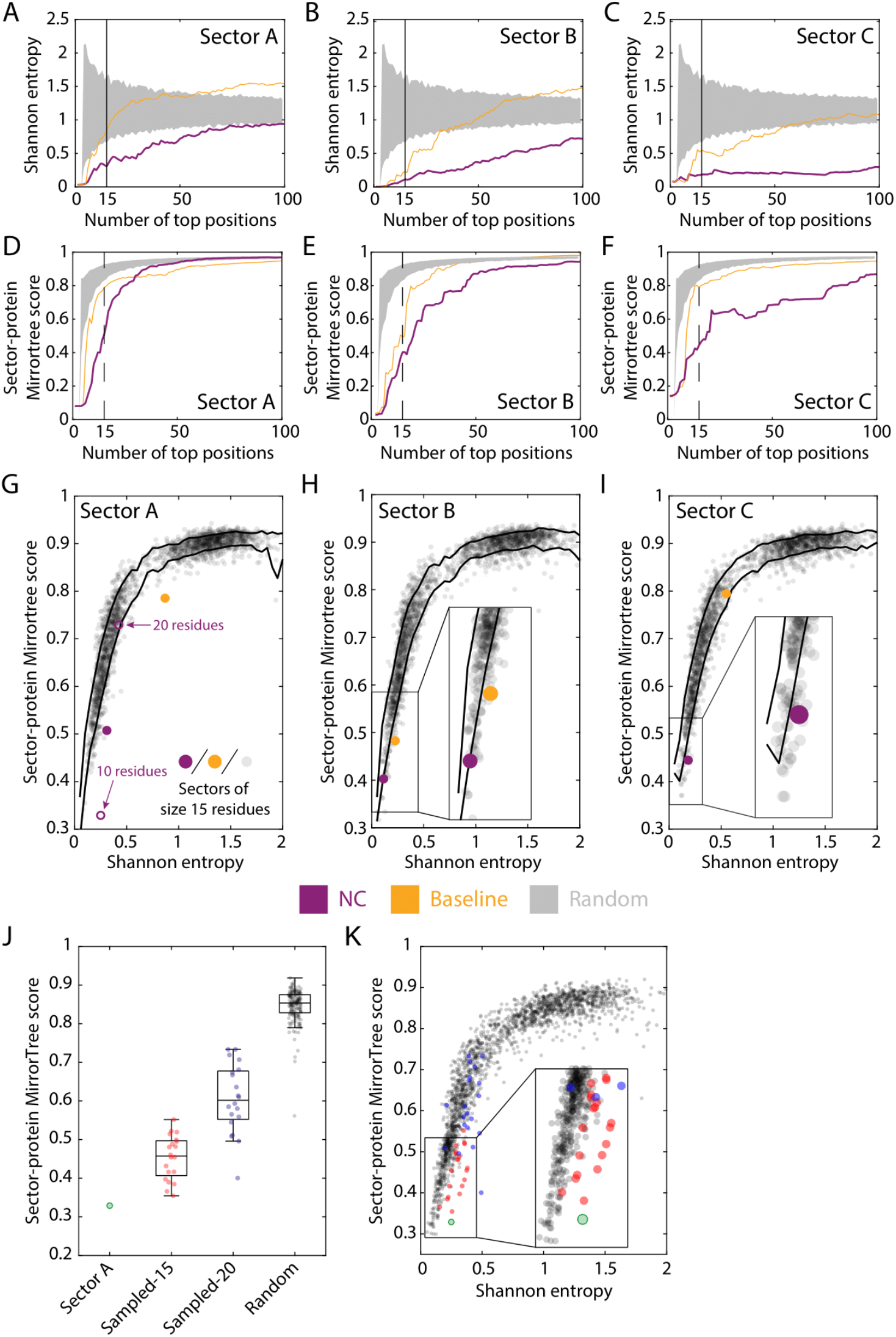
Sector-protein MirrorTree scores of residue groups are correlated with entropy, but NC sectors have lower MirrorTree scores than expected from entropy alone. A-C) The Shannon entropy of MreB NC sectors A-C (Fig. 3) across size cutoffs is lower than that of the corresponding baseline sectors, indicating that NC selects more conserved residues (albeit entropy is still higher than the cutoff of <0.1 for typically being considered highly conserved). Gray regions represent the entropy of a randomly selected group of residues of the same size. D-F) MirrorTree scores are lower for the NC sectors than for the corresponding baseline sectors. Gray regions represent the MirrorTree scores of a randomly selected group of residues of the same size. G-I) The MirrorTree scores of sectors A-C (filled gold and purple circles) and of random groups of 15 residues (gray). Although MirrorTree score is linked to entropy, NC sectors A and C have MirrorTree scores significantly lower than expected based on entropy alone. In (G), the open purple circles denote the versions of sector A with 10 and 20 residues. Black curves indicate ±1 standard deviation from the mean MirrorTree score for a given entropy. J) The 10-residue version of NC sector A has lower MirrorTree score than sets of 10 residues selected from the 15- and 20-residue versions of the same sector, which are lower than those of random groups of 10 residues. The central mark indicates the median, and the bottom and top edges of the box indicate the 25^th^ and 75^th^ percentiles, respectively. The whiskers extend to the most extreme data points not considered outliers. K) The 10-residue version of NC sector A has a lower MirrorTree score than 10-residue subsets of the 15- and 20-residue versions of the same sector with similar entropy. Same data as in (J). Thus, the 10-residue sector represents a “core” of the most highly coevolving residues.

MirrorTree scores of NC sectors were also generally lower than those of baseline sectors (Fig. 5D-F). To investigate the dependence of sector-protein MirrorTree scores on entropy, we computed MirrorTree scores for thousands of random groups of the same size as the sector (15 residues), biasing sampling using a Monte Carlo algorithm to obtain a wide range of mean entropies; each random group was selected from residues that did not overlap with the sector. For mean entropy ≾1, MirrorTree scores were strongly dependent on entropy (Fig. 5G-I). Thus, the low MirrorTree scores of the NC sectors were due in part to their low entropy. Nonetheless, the MirrorTree score of NC sector A was significantly lower than those of random groups with the same mean entropy (*z*-score −3.5); the entropy of sector B was so low, presumably due to the high conservation of the ATP-binding pocket (Fig. 3H, S4), that it was challenging to obtain random groups that were not largely overlapping.

Since NC sector A displayed the greatest reduction in MirrorTree score relative to random groups of the same mean entropy, we focused on this sector to investigate the dependence of the sector-protein MirrorTree score on sector size. As the cutoff was increased to include more residues, the MirrorTree score increased (Fig. 5D). To disentangle whether this increase was due directly to the increase in size or to the inclusion of residues that are more phylogenetically similar to the protein, we compared the 10-residue version of sector A (Fig. 5G) with randomly selected groups of 10 residues from 15- and 20-residue versions of sector A, as well as the entire protein. The mean MirrorTree score increased as the size of the sampling group increased (Fig. 5J), even for groups with similar entropy as the 10-residue sector (Fig. 5K). Moreover, 15-residue versions of sectors B and C had similar entropy (Fig. 5B,C); hence, an approach driven by entropy alone would not have divided these spatially separated clusters. Thus, the strength of a residue’s association in a sector of highly coevolving residues is associated with more phylogenetic distinction from the rest of the protein than can be explained by entropy alone.

### Phylogenetic similarity highlights overlapping sectors

The core residues of some MreB NC eigenvectors sometimes had high coefficients in multiple eigenvectors (Fig. S5), suggesting that we should consider the union of the sectors as a functional unit. To rationally identify sectors that should be merged, we again exploited phylogenetic similarity by calculating MirrorTree correlation coefficients from comparisons between pairs of sectors (Fig. 6A). MreB NC sectors A-C (Fig. 3) exhibited low sector-sector MirrorTree scores with each other and with random groups (Fig. 6B), as expected since they have low sector-protein MirrorTree scores (Fig. 6B). By contrast, the random groups had MirrorTree scores close to 1 (Fig. 6B). NC sectors were also more phylogenetically distinct from each other than baseline sectors (Fig. 6C). These data suggest that the NC sectors were selected by evolutionary pressures that led to distinct functions.

**Figure 6:**
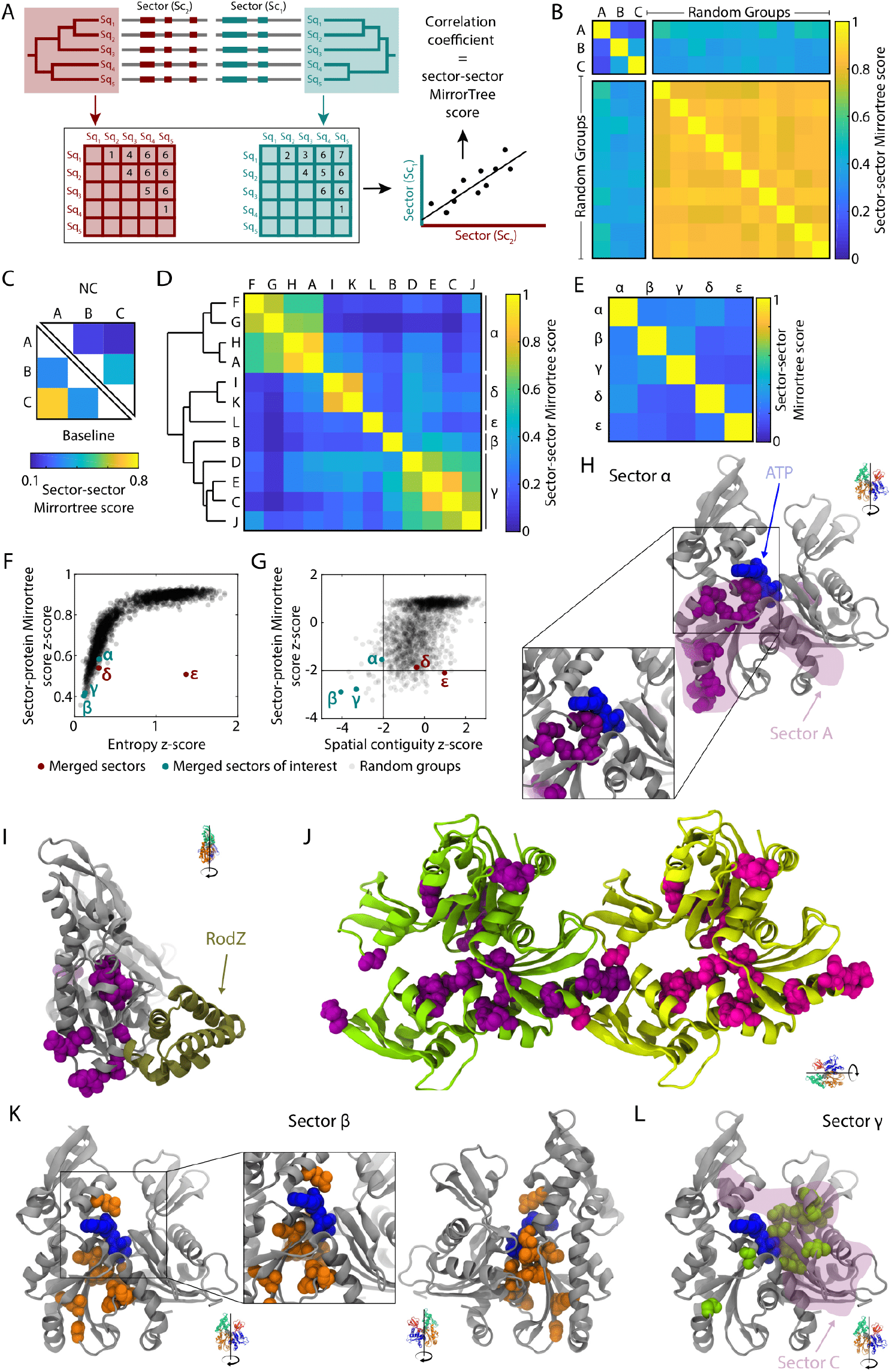
MreB NC sectors are generally phylogenetically distinct, and those with phylogenetic overlap collectively overlap with functionally important regions. A) Repurposing the MirrorTree algorithm to measure the phylogenetic similarity between sectors. B) MreB NC sectors A, B, and C exhibited low sector-sector MirrorTree scores with each other, but high values with random groups of 15 residues (which also exhibited high MirrorTree scores with each other). C) NC sectors A-C have lower sector-sector MirrorTree scores with each other than baseline sectors A-C with each other, indicating that they are more phylogenetically distinct. D) Hierarchical clustering of MreB NC sectors A-L (Fig. 4C) based on sector-sector MirrorTree profiles suggests five distinct mega-sectors. E-G) The MreB mega-sectors defined by the sum of the clustered eigenvectors exhibited low sector-sector MirrorTree scores with each other (E) as well as low sector-protein MirrorTree scores (F). Mega-sectors α, *β*, and γ (similar to sectors A-C) exhibited high spatial contiguity (*z*-score<-2). H,I) Mega-sector α was more spatially contiguous than sector A (shaded purple region) (H), and contained residues around the interface with MreB’s binding partner RodZ (I). J) The 25-residue version of mega-sector α connects the pointed and barbed ends of each subunit in a protofilament. K) Mega-sector *β* (identical to sector B) surrounds the ATP binding pocket. L) Mega-sector γ is more spatially contiguous than sector C (shaded purple region).

Of all sectors that had a MirrorTree *z-*score or a pairwise distance *z*-score<-2 (sectors A-L, Fig. 4C), several pairs had a high sector-sector MirrorTree score. Hierarchical clustering of the sectors based on their sector-sector MirrorTree profiles led to the identification of five obvious “mega-sectors” from the sum of the clustered eigenvectors (Methods), which we denote α, β, γ, δ, and ε(α, β, and γ contain sectors A, B, and C, respectively) (Fig. 6D). The mega-sectors exhibited low sector-sector MirrorTree scores (Fig. 6E), and α, β, and γ had both low sector-protein MirrorTree scores (Fig. 6F, G) and low spatial contiguity *z*-scores (Fig. 6G). The 15-residue version of mega-sector α more compact than the 15-residue version of A (Fig. 6H), and it contained residues that interact with RodZ (Fig. 6I), an MreB binding partner that modulates MreB filament nucleation (44) and curvature (45). Notably, the regions of the 25-residue version of mega-sector α at the barbed and pointed ends of the MreB subunit interact with each other in a polymerized MreB filament (Fig. 6J), reinforcing the spatial contiguity of the mega-sector. Mega-sector *β* was identical to sector B, surrounding the ATP-binding pocket (Fig. 6K). As with α and A, the 15-residue version of mega-sector γ was more compact than the 15-residue version of sector C (Fig. 6L), indicating that clustering based on MirrorTree scores increases the spatial contiguity of sectors.

### NC identifies sectors that are not apparent from the full coevolution matrix

To determine whether our findings about the properties of NC sectors applied to other proteins, we performed similar sector calculations for enolase (the metalloenzyme responsible for conversion of 2-phosphoglycerate to phosphoenolpyruvate during glycolysis (46); Fig. 7A-C), the carbohydrate-processing enzyme glucose-6-phosphate dehydrogenase (G6PD (47); Fig. 7D-F), and mitogen-activated protein kinase 1 (MAPK1) (48, 49) (Fig. 7G-K). In each case, NC produced sectors with lower background noise and higher spatial contiguity than baseline sectors.

**Figure 7:**
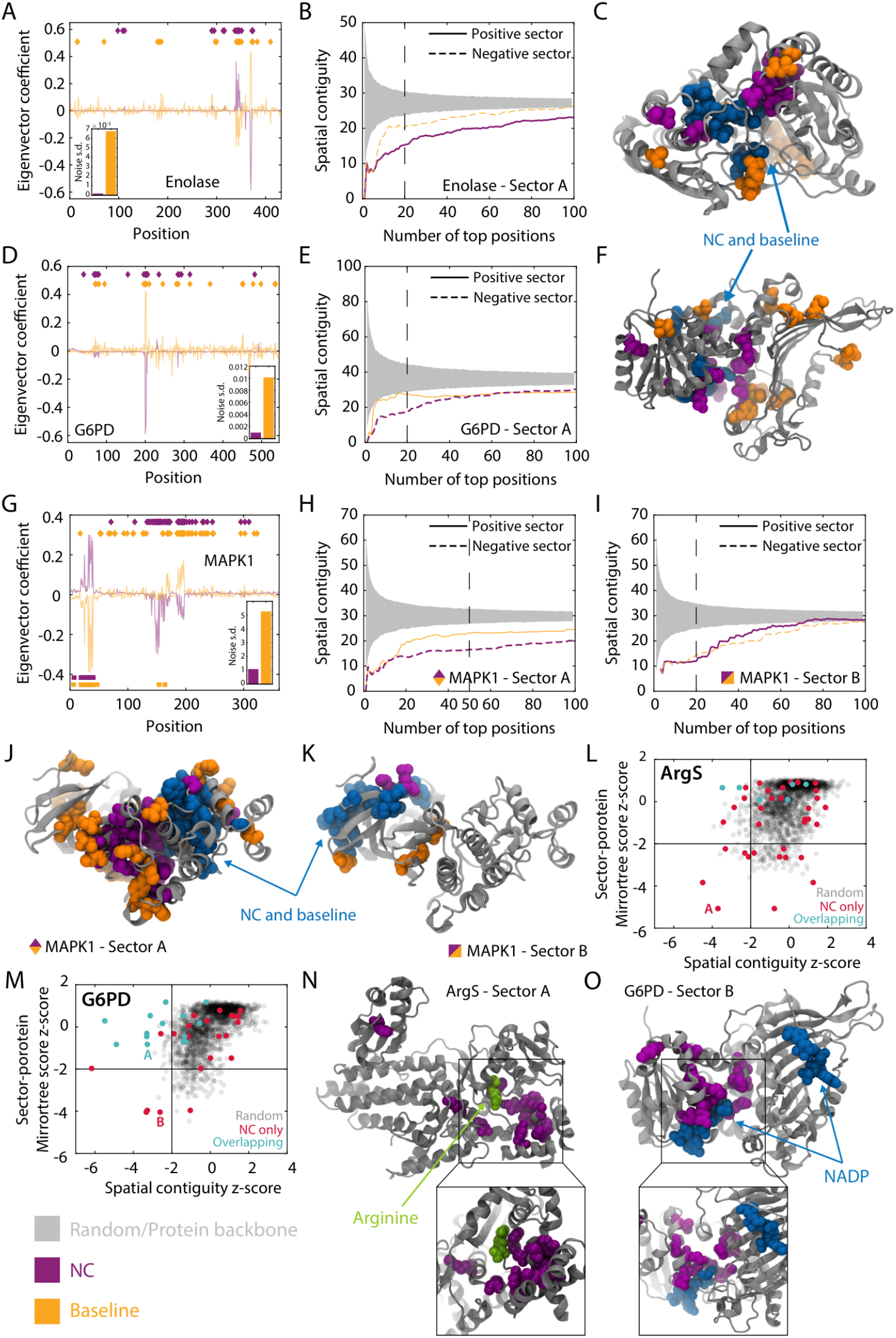
NC eigenvectors generally improve sector prediction across proteins, and enable identification of sectors that are not detectable using the baseline method. A,D,G) NC eigenvectors for enolase (A), G6PD (D), and MAPK1 (G) exhibit lower background noise than the corresponding baseline (NMI with APC) eigenvectors. B,E,H,I) The appropriate NC sectors (positive or negative values of the eigenvector) associated with the eigenvectors in (A,D,G) are more spatially contiguous across size cutoffs than the baseline sectors. Note that the MAPK1 eigenvector was split into a positive sector (H) and a negative sector (I). C,F) The 15-residue versions of the sectors in (B,E) on the crystal structures of enolase (C) and G6PD (F) illustrate the more compact nature of the NC sectors as compared with the baseline sectors. J,K) The 50- and 20-residue versions of the NC sectors in (H,I) are more spatially compact on the structure than the corresponding baseline sectors, and occupy distinct parts of the protein. L-O) For ArgS (L) and G6PD (M), certain high-eigenvalue NC sectors had no obvious baseline counterpart. These NC sectors had low MirrorTree and spatial contiguity *z*-scores (L,M), and 15-residue versions occupied spatially compact regions around ligands (arginine in (N), NADP in (O)) on the structure (N,O). Thus, NC enables the detection of sectors that are otherwise hidden.

Most of the MreB NC eigenvectors had strong signal for either positive or negative coefficients, but not both (Fig. 3A-C). By contrast, one of the large-eigenvalue NC eigenvectors for MAPK1 had groups of residues with both very positive and very negative coefficients (Fig. 7G); these residues were located in distinct regions of the protein (Fig. 7J,K). As validation for splitting the NC eigenvector into two sectors, the sector-sector MirrorTree score (0.44) indicated that they are phylogenetically distinct; moreover, the sector-sector MirrorTree score of the corresponding baseline sectors was higher (0.71). Thus, NC eigenvectors can be interpreted as two phylogenetically distinct sectors based on coefficient signs.

In addition to improving sector predictions by reducing background variation, we were interested in determining whether NC is able to identify sectors that the full coevolution matrix misses altogether. For the arginine tRNA ligase ArgS (50) and G6PD, the sector with the most negative MirrorTree *z*-score had nearly the lowest spatial contiguity *z*-scores (Fig. 7L,M) and no clear counterpart in any of the baseline eigenvectors (Methods). For ArgS, the NC sector was spatially localized around the arginine binding site (Fig. 7N). For G6PD, the NC sector was adjacent to one of the two NADPs that bind to the protein (Fig. 7O). Thus, the NC correction reveals some sectors that are missed by the baseline method.

### NC sectors are enriched in damaging mutations

To test the functional significance of NC sectors, we sought experimental datasets with quantitative measurements of the consequences of mutations across a protein of interest. Recent studies have pioneered the use of deep mutational scanning to systematically generate and quantify the phenotypic or fitness effects of a large number of individual mutations spanning entire domains or protein (29, 51–53), providing new insights into structure-function relationships. Thus, we asked whether NC sectors were enriched in residues for which mutation altered protein function and/or fitness.

The Ras superfamily of membrane-associated small G-proteins is highly conserved and controls a broad range of cellular processes (54), has inactive and active states that are regulated by a GTPase-activated protein (55), and has been implicated in cancer (56). A recent deep mutational scanning study engineered plasmids to express mutant versions of human H-Ras as well as the Ras-binding domain of human C-Raf (Raf-RBD) in *Escherichia coli* (57), such that the binding of Ras·GTP to Raf-RBD led to transcription of a chloramphenicol-resistance cassette. Thus, the binding efficacy of the Ras variant was directly correlated with cellular growth rate. The effect of Ras mutations on fitness was quantified by the logarithm of the enrichment of variants in the chloramphenicol-selected versus the starting population, relative to wild-type. The distribution of fitness effects was centered around zero, although there were some positions with mutations that displayed significant functional effects (57).

To determine whether fitness-altering mutants in H-Ras are enriched at positions identified by coevolution, we identified two high-eigenvalue sectors with obvious corresponding baseline sectors. As in our previous analyses (Fig. 3A-C, 7A,D,G), aside from the highly coevolving residues, the NC sectors had much lower noise than the baseline sectors (Fig. 8A,B). The residues in the two NC sectors were non-overlapping, and in both cases appeared to be concentrated in regions with low minimum relative enrichment (Fig. 8C,D). Across cutoffs that defined sectors of various sizes, we computed the minimum and maximum relative enrichment (representing deactivation and activation, respectively) over all amino acid mutations for each position in the NC/baseline sectors as well as for the residues with the lowest entropy, and compared to the distribution over all residues. As expected, the residues with lowest entropy consistently predicted significantly more negative minimum relative enrichment than random sets of residues (Fig. 8E,F). The mean minimum relative enrichment in NC and baseline versions of sector A was also significantly more negative than random residues, with the NC sector outperforming the baseline sector and achieving similar enrichment values to the lowest-entropy residues (Fig. 8E). NC sector B also exhibited mean minimum relative enrichment significantly lower than random, by contrast to the baseline sector (Fig. 8F). Thus, sectors A and B are more enriched for residues whose mutation has the most potential for reducing fitness using NC versus baseline. The maximum relative enrichment was highly similar for sectors and the protein overall (Fig. S6A,B), suggesting that NC and baseline sectors are enriched for residues with the potential for deactivating rather than activating mutations in the case of H-Ras. Thus, NC sectors separate residues based on the maximum impact of mutations at these positions.

**Figure 8:**
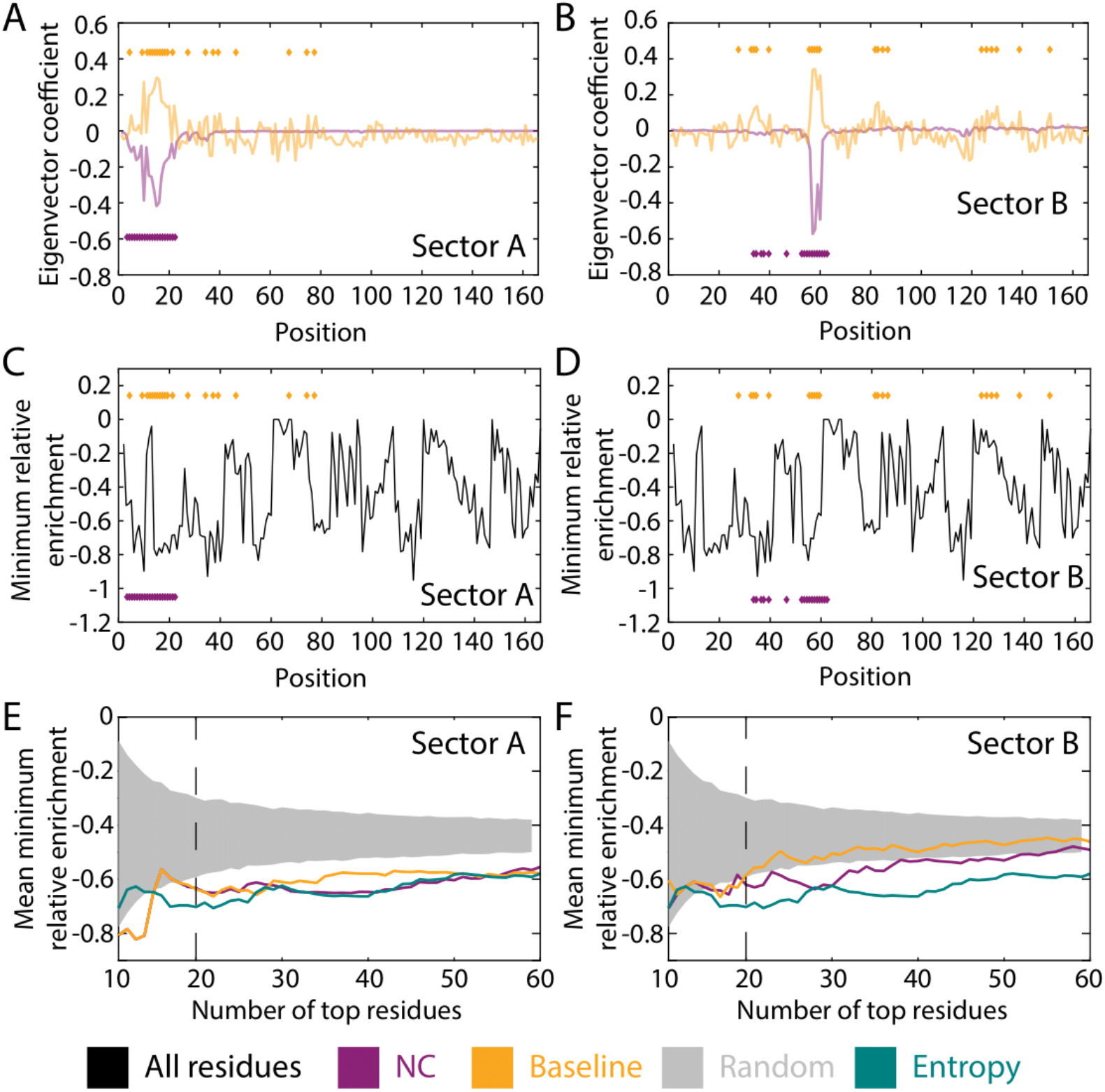
NC sectors predict deactivating mutations in H-Ras. A,B) NC predicts two eigenvectors with much lower background noise than the baseline counterparts. The purple and gold diamonds represent the locations of residues in sectors of size 20. C,D) Fitness data from a screen of binding efficacy of H-Ras to Raf-RBD (57). Shown is the minimum enrichment over all mutations at each position (thus representing maximum deactivation). The purple and gold diamonds represent the locations of residues in sectors of size 20. E,F) Across most sector size cutoffs, the mean minimum relative enrichment was significantly lower than random (gray) for NC sectors A and B and comparable that of the residues with the lowest entropy (teal). NC sectors also outperformed their baseline counterparts.

## Discussion

Many existing coevolution methods build on correlation or mutual information, sometimes employing ad-hoc corrections to partially remove the effects of entropy and phylogeny. Our NC method harnesses phylogenetic distance between sequences as a novel dimension in the measurement of protein coevolution, in order to increase understanding of the functional relationships between amino acids in a protein. In particular, here we demonstrated that coevolution can occur on multiple phylogenetic timescales within a single protein. While the factors that determine whether pairs of positions coevolve on short or long timescales are unknown, future studies using NC to interrogate the specific biochemical functions of protein sectors may reveal general patterns across diverse proteins. One interpretation of the variable contribution of coevolution across phylogenetic distance within a single protein (Fig. 1C) is that the frequency of mutation for coevolving residues within an NC sector is linked to the timescale of change for the corresponding selective pressure on that sector. For example, a sector that determines protein thermostability would be predicted to coevolve on a timescale commensurate with the frequency of changes in environmental temperature, whether these changes occur over long (e.g. glaciation and interglacial cycles of 100,000 years) or shorter (e.g. Atlantic multidecadal oscillations) timescales.

Importantly, NC and our repurposed MirrorTree methods are complementary to most covariation metrics, and hence can enhance existing bioinformatics tools by defining a phylogenetic dimension of coevolution and allowing focus on functional signal. We anticipate that our approach will enable application of coevolution-based methods across a much broader class of proteins, including those for which the set of sequences is limited in number (Fig. 2) and/or for which the available homologous sequences are biased to a particular segment of the phylogenetic tree (Fig. 1B). In particular, application to the growing database of human exome sequences (58) may improve identification of rare disease-causing mutations. NC may also enhance protein engineering tools by highlighting targets for directed evolution. As we have demonstrated, NC expands our ability to detect functional relationships between residues within proteins and to determine the links between protein evolution and adaptation. In concert with deep mutational scanning and other comprehensive functional screens (59), NC and MirrorTree should provide deeper insight into the specific selective pressures under which proteins have evolved.

The predominant application of coevolution so far has been structure prediction, from using top DCA-predicted contacts as constraints (4) to employing DCA model parameters as input training features for deep neural networks that seek to predict spatial distances between amino acids (60). Here, we have shown that NC can improve contact prediction by DCA. Moreover, the detection and interpretation of sectors as functional units within proteins has been a growing research focus, particularly with respect to the evolutionary origins of sectors. A recent theoretical study demonstrated that selection acting on a functional property can give rise to a sector (28). Here, we showed that NC better resolves sectors than baseline by reducing background noise (Fig. 3A-C), leading to sectors with higher spatial contiguity (Fig. 3D-L) and lower MirrorTree scores (Fig. 4B). Low MirrorTree scores reveal that residues within sectors have a different evolutionary history from the rest of the protein, due to both entropy-dependent and entropy-independent differences (Fig. 5). MirrorTree scores can further be used to evaluate NC predictions in the absence of a known structure. Motivated by the original design purpose of MirrorTree, we note that scores between sectors of two proteins could be used to identify protein-protein interactions—potentially between hosts and microbes—due to the improved performance of NC when the sampling of sequences is shallow (Fig. 2C).

Our observation that NC sectors, and moreover their cores, have high spatial contiguity and low MirrorTree scores (Fig. 5J) supports the inferred link between coevolution and spatial contiguity, and suggests that NC can help to guide experiments toward the residues of highest importance for a sector’s function (Fig. 8). Beyond the improvements from lowering background signal, NC also predicts sectors that are otherwise difficult to detect (Fig. 7L-O), thus highlighting its value. In addition, some studies have demonstrated other applications such as protein engineering (19) and variant interpretation (14). Improved detection of functional coevolution could even help to refine MSA algorithms, which are ultimately a limiting factor in the detection of coevolution. Our results suggest that the utility of coevolution as a signal for protein science can be substantially improved by NC, opening new windows for broadly understanding (and perhaps ultimately engineering) protein structure-function relationships.

## Methods and Materials

### MSA construction

MSAs were constructed with BLAST (61) to identify up to 10,000 closest sequences to a reference sequence, using the RefSeq database (62). Sequences were aligned with Clustal Omega (63). Sequences with a Jukes-Cantor distance >1 from the reference sequence were pruned. Redundant sequences and positions with >25% gaps were removed. Any remaining gaps were filled with the closest amino acid from the closest sequence in terms of Jukes-Cantor distance.

### Calculation of the expected value of inter-clade covariation

For our analyses, we define a pair of sequences to be within the same clade if the phylogenetic distance is below a Jukes-Cantor distance *d*. The phylogenetic distance is measured with respect to the aligned protein sequence (Table S1). We sought to measure the expected value of residue-residue covariation due solely to the comparison of sequences between clades, which we refer to as the inter-clade covariation *C*_*S*>*d*_. Below, we describe and compare measurement of the expected value of inter-clade covariation in Eq. 1 of the main text both by approximation via bootstrapping and analytically.

#### Bootstrapping

In this approximate method, we bootstrap the original MSA: for every position, we replace the amino acid with the identity of the same position from a random sequence in the same clade. For example, in Fig. 1Avi we show two positions in an MSA, colored by their clade membership for a given phylogenetic distance *d*. Note that the first position is never a glutamine in the orange clade and is never a threonine in the white clade. Similarly, the second position is never a serine in the orange clade and is never an arginine in the white clade. The bootstrapped MSAs resample within clades, so as to not change the phylogenetic structure of the MSA at distances >*d*; thus, the first position in the bootstrapped MSAs still does not contain a glutamine, etc. The covariation measured from each of the bootstrapped MSAs is averaged to obtain the matrix expected under the hypothesis that there is no coupling between positions within the same clade. The bootstrapping method can be applied for any coevolution heuristic.

#### Analytical method

To derive an analytical solution in place of bootstrapping the NMI metric, we rephrased our aim as calculating the expected value of covariation between two positions under the assumption that the two positions are independent within a clade.

Consider the Shannon entropy for position *i*:

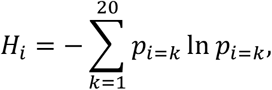

where *p*_*i*=*k*_ is the probability of finding amino acid *k* at position *i* . The marginal probabilities of positions *i* and *j* taking on a particular value in a bootstrapped MSA do not change on average. However, the joint entropy, which relies on the joint probability, will change:

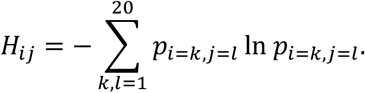

We seek an expression for the joint entropy that captures the assumption that positions *i* and *j* are independent within clades. Since the joint probability of independent variables is the product of the individual probabilities, we are left with calculating the sum of probabilities from each clade *c*, weighted by the number of sequences *n*_*c*_ in each clade:

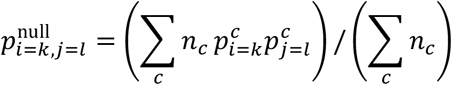

where 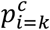 is the marginal probability of finding amino acid *k* within clade *c* at position *i*.

A comparison of the bootstrapped and analytical methods for calculating NC for the yeast actin protein is shown in Fig. S2.

#### Estimating the statistical significance of nested coevolution

The expectation value of our nested coevolution background model is described above analytically only for normalized mutual information; other coevolution metrics do not have a known closed form analytical solution, so we rely on bootstrapping to estimate the expected value. Bootstrapping offers the additional advantage of providing an estimate of the statistical significance of the observed raw coevolution signal by measuring what fraction of bootstrapped MSAs achieve equal or greater coevolution values. The accuracy of the significance estimate is limited by the number of bootstrap measurements, since the maximum resolution is the reciprocal of the number of bootstraps performed. Using hundreds of bootstraps, we compared significance estimates with the absolute difference between the total and inter-clade covariation. These values were highly correlated (Spearman’s *ρ* = 0.95, Fig. S2B), indicating that either the bootstrapping or analytical method of computing NC provides a surrogate for the significance of the observation.

### Structural contact prediction

Real structural contacts were determined by calculating the distance between the alpha carbons of every pair of residues in the protein based on a crystal structure (Table S1). All other atoms, including hydrogen atoms, were disregarded. To predict structural contacts, we used mean-field DCA, and the value of the pseudocount is 0.5, and sequences closer than 0.3 Hamming distance are reweighted (4, 22).

### Generation of NC sectors

The output of NC is *n*_*d*_ *p*-by-*p* matrices (Fig. 1C), where *n*_*d*_ is the number of phylogenetic windows and *p* is the number of amino acids in the protein. These *n*_*d*_ matrices are concatenated to obtain a supermatrix of dimension *pn*_*d*_-by-*p* (Fig. S7). Principal component analysis using eigenvalue decomposition or singular value decomposition is performed on the super matrix (thus avoiding the need to choose one value of the cutoff distance *d*), with *pn*_*d*_ observations and *p* features. The eigenvectors are ordered highest to lowest according to their associated eigenvalues. Each eigenvector is of length *p*, where the *i*^th^ coefficient corresponds to the importance of the *i*^th^ amino acid in explaining the variation in the direction of the respective eigenvector.

To extract the specific amino acids that are most responsible for explaining the variation in a particular eigenvector, we identify the positions with the most positive or most negative coefficients and define these groups of residues as two sectors. Sectors that have <4 amino acids are ignored for downstream analysis.

NC and baseline sectors were paired if the dot product of the corresponding eigenvector was >0.6.

### Calculating the spatial contiguity of a sector

To quantify spatial contiguity, we calculate the mean distance between the alpha carbon atoms of each pair of residues in the sector in the crystal structure.

### Adaptation of the MirrorTree algorithm

Mirrortree was originally developed to predict protein-protein interactions based on the similarity of phylogenetic trees (37). In brief, MSAs are calculated using protein sequences from the same list of organisms for two proteins. For each MSA, the matrix of pairwise Jukes-Cantor distances is calculated. The MirrorTree score is the Pearson correlation coefficient of these two distance matrices. A high correlation indicates that the two proteins have similar phylogenies and thus are likely to have experienced similar functional selection. We adapted this method to compare the phylogenetic similarity of protein sectors with the entire protein (Fig. 4A) or other sectors (Fig. 6A). To compute sector-protein and sector-sector MirrorTree scores, filtered MSAs were created focusing on the positions of a given sector.

Biased sampling of random sectors was accomplished via weighting of residues according to their entropy.

### Calculation of megasectors

Sets of sectors to be merged into megasectors were determined from hierarchical clustering based on sector-sector MirrorTree scores. Merging was accomplished by adding the corresponding eigenvectors after multiplying each sector by +1 or −1 corresponding to whether a positive or negative sector, respectively, was being merged. The summed vector was then analyzed as if it were an eigenvector in order to define megasectors at various size cutoffs.

## Acknowledgments

The authors thank the Huang lab for useful discussions. This work was supported by Stanford Graduate Fellowships (to A.C. and E.A.), a Gerald J. Lieberman Fellowship (to A.C.), grant 851173 from the European Research Council under the European Union’s Horizon 2020 research and innovation programme (to A.-F.B.), NSF CAREER Award MCB-1149328 (to K.C.H.), and the Allen Discovery Center at Stanford on Systems Modeling of Infection (to K.C.H.). K.C.H. is a Chan Zuckerberg Biohub Investigator.

