## Supplemental Figures/Table for "Extracting the phylogenetic dimension of coevolution reveals hidden functional signal"

### Supplementary Figures

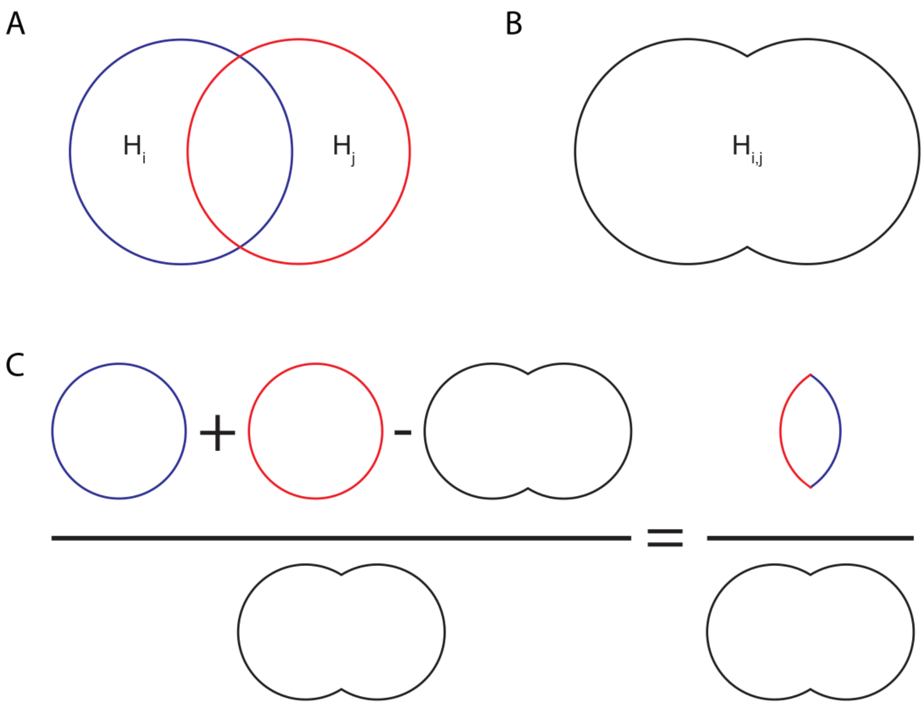


**Figure S1: Pictorial representation of information-theoretic metric of residue-residue coupling.**

A) Protein positions *i* and *j* that take on multiple amino-acid identities in an MSA have non-zero entropies, represented by the areas of $H_i$ and $H_j$. The two circles are overlapping because knowing the identity of position *i* provides some information about the identity of position *j*.

B) The union of the space occupied by positions *i* and *j* is the joint entropy.

C) The normalized mutual information metric of coevolution (Eq. 1) represents the sum of the entropies in (A) minus the joint entropy in (B), normalized by the joint entropy in (B).

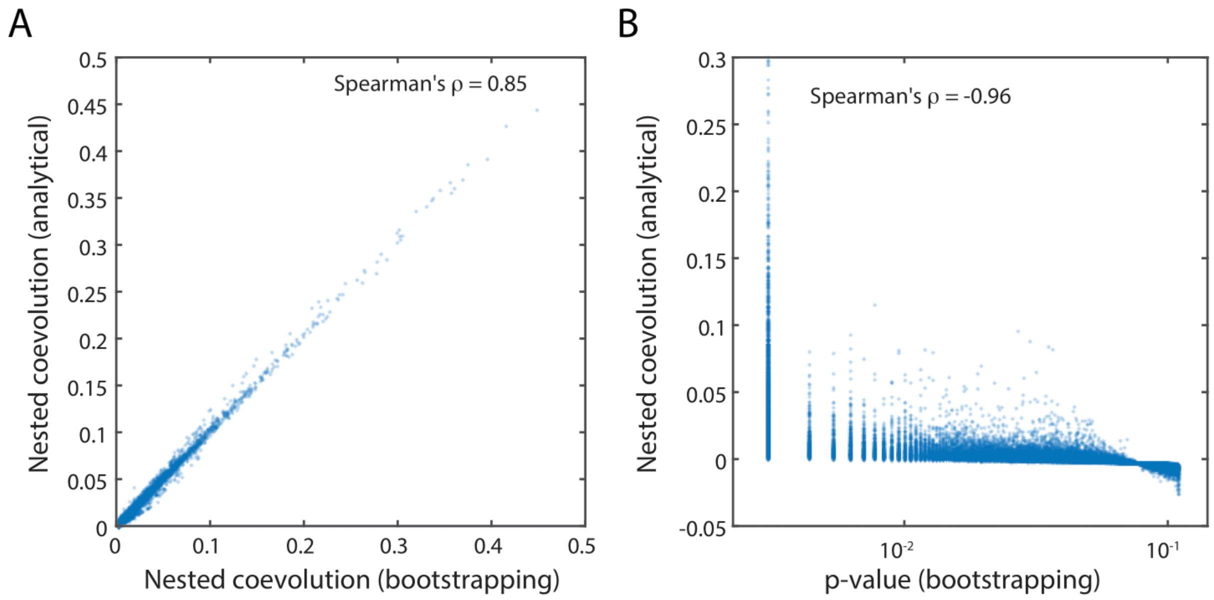


**Figure S2: Relationship between magnitude and significance of NC values.**

A) NC measurements by bootstrapping and analytical methods (Methods) are highly correlated.

B) The significance of NC, estimated by bootstrapping (Methods), is highly correlated with its magnitude, demonstrating that magnitude of NC signal is a surrogate for its significance.

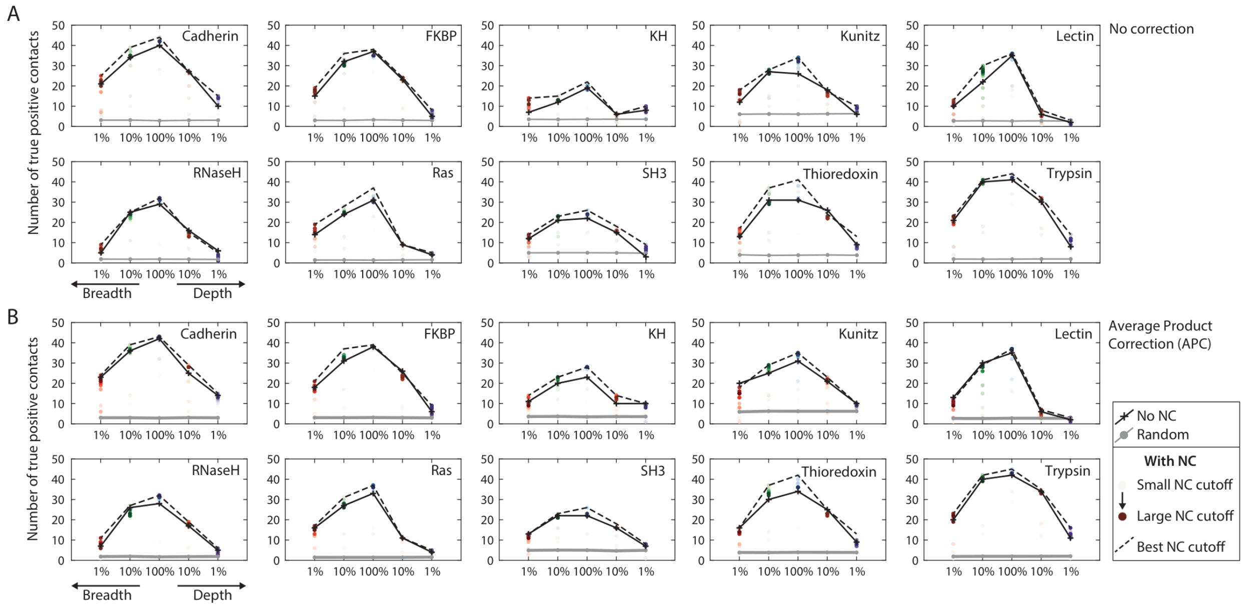


**Figure S3: NC improves structural true-positive contact prediction over DI + APC.**

NC delivers a net improvement in recovering structural information by DCA, which is stronger for breadth-sampled MSAs than depth-sampled MSAs, compared to baseline (in this case DI with APC). NC recovers breadth better than depth, in absolute terms highlighting the importance of having breath in an MSA. NC seems to provide approximately the same fold-change in improvement across subsampling degrees.

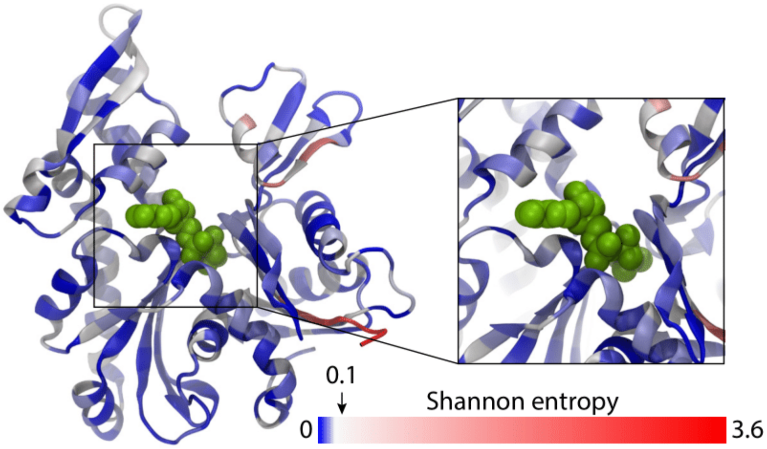


**Figure S4: The ATP binding pocket of MreB is composed of highly conserved residues.**

Amino acids are colored by their Shannon entropy, with blue indicating values <0.1. The ATP molecule is shown in green.

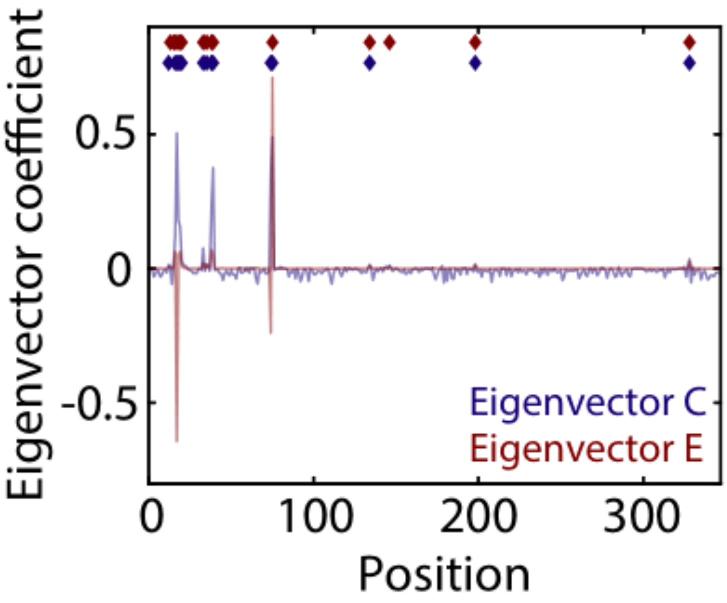


**Figure S5: Similar eigenvectors are merged to create mega-sectors.** The positive potions of eigenvectors C and E define sectors C and E, respectively. These eigenvectors are combined to create a mega-sector.

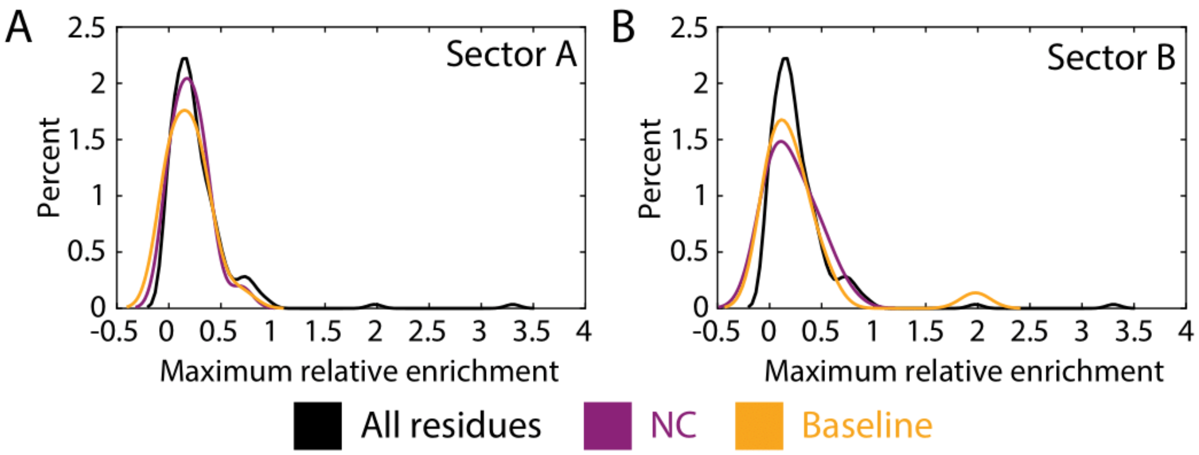


**Figure S6: The maximum relative enrichment for H-Ras sectors is similar to the enrichment for the protein overall.** The maximum enrichment profile for sectors A (A) and B (B) are similar to that of the overall protein (black) for both NC and baseline.

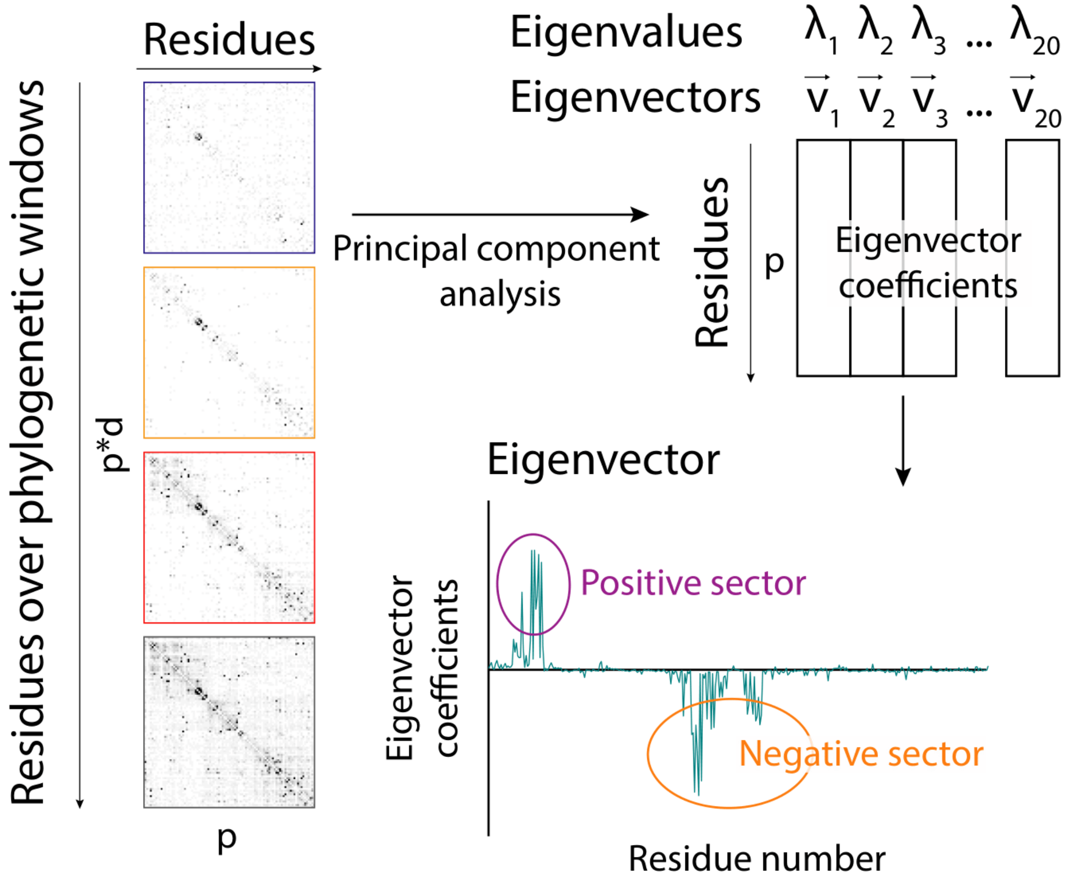


**Figure S7: Schematic showing sector determination.** The NC sectors are concatenated, principal component analysis is performed, and the eigenvectors associated with the highest eigenvalues are used to define positive and negative sectors.

**Supplementary Table**

| Protein | Uniprot ID | NCBI accession number | PDB ID |
|---|---|---|---|
| Cadherin | I3LUS1_PIG | - | 2O72 |
| Enolase | - | NP_417259 | 5OHG |
| FKBP-C | A0A0D2TZB3_GOSRA | - | 1R9H |
| G6PD | - | NP_000393 | 2BH9 |
| H-Ras | - | 1BKD_R | 1BKD |
| KH-1 | B4MJF9_DROWI | - | 1WVN |
| Kunitz | F6WEH0_ORNAN | - | 5PTI |
| Lectin-C | H0YSP5_TAEGU | - | 2IT6 |
| MAPK1 | - | NP_620407 | 6FN5 |
| MreB | - | NP_417717 | 2WUS, 1JCG, 4CZE |
| Ras | A0A024W598_PLAFA | - | 5P21 |
| RNase H | R9T5K3_9EURY | - | 1F21 |
| SH3 | A0A0K0FYI6_9BILA | - | 2HDA |

| | | | |
|---|---|---|---|
| Thioredoxin | H0ZH31_TAEGU | - | 1RQM |
| Trypsin | I3M2H8_ICTTR | - | 3TGI |

**Table S1: Reference sequences and PDB structures.**
